# Unveiling consistency in flexibility: the role of reward and cognitive control in moral decisions

**DOI:** 10.1101/2023.06.17.545416

**Authors:** Xinyi Julia Xu, Guochun Yang, Jiamin Huang, Ruien Wang, Haiyan Wu

## Abstract

Moral decisions are multifaceted with two essential aspects, flexibility and consistency. However, the interaction between these two and the underlying mechanisms is rarely studied. Here, we combined mouse-tracking and functional magnetic resonance imaging (fMRI) together in a value-based moral decision task, which allows us to quantify accumulative history responses as self-consistency. Using a multi-attribute time-dependent drift-diffusion model (tDDM), we disentangled the role of consistency and self-interest, highlighting the dominant role of cognitive-control-related regions. The drift rate of self-consistency was directly associated with the brain activity responsible for cognitive control, while the relationship between the reward and the activity of the related brain regions was mediated by the mouse-tracking index area under the curve(AUC). Dorsolateral prefrontal cortex was revealed as a hub connecting prACC and ventral striatum, whose functional connectivity was correlated with consistency drift rate and reward drift rate respectively. Furthermore, decision flexibility was quantified by choice entropy, which links to mouse tracking indices and the activity of cognitive -control related regions. Together, our study uncovers the interplay between self-consistency and reward in behavior, and highlights the key role of cognitive control in modulating these two attributes, thereby deepening our understanding of consistency and flexibility in moral decisions.

## Introduction

Flexibility and consistency are two opposing forces that shape our decisions^1^. On one hand, people adapt their decision-making strategies to fit different social contexts (i.e., **flexibility**). For example, if they happen to have lunch with vegetarians, they may prefer selecting vegetable-based dishes, whereas when dining with seafood enthusiasts, they could lean towards opting for shrimp-based meals. When it comes to morality, moral flexibility refers to the beliefs guiding people’s behavior depending largely on context, even when people really intend to be moral^2^. On the other hand, there is also a sense of **consistency** that permeates our decisions. People tend to keep consistent responses and avoid contradictory arguments under repeated situations, which is referred to as internal consistency^3,4^. The aspects of flexibility and consistency are intricately intertwined, influencing and interacting with each other in moral decisions. Within an individual, moral decision-making can be subject to fluctuations based on the individual’s emotional state, cognitive biases, and subjective interpretation of ethical principles. These volatile factors can lead to inconsistencies and unpredictability in moral decision-making, making inconsistent choices.

The pursuit of self-interest, one main factor that leads to self-serving decisions, is a key feature of human decision-making^5,6^. For example, although honest responses might require fewer cognitive control resources and hence a default strategy, dishonesty may be the faster response when self-interest overrides the preference for honesty^7^. One characteristic of self-serving decisions is that humans are more likely to make a selfish choice as the reward or self-interest increases. Another factor plays a significant role in moral decisions is self-image^8^. Previous literature on moral decision-making illustrates that moral versus immoral decisions are driven by both rewards and self-image^7,9^. In the case of honest/dishonest, seeking rewards may lead to moral norm conflict; thus, people sometimes perform dishonest acts consciously and deliberately by trading off between the expected external benefit and the moral cost of the dishonest acts^10,11^. Over the past few decades, studies have demonstrated that complex moral choices can be predicted by the utility of combining these two considerations (expected external benefit and the moral cost of the dishonest acts)^9,12^. However, few studies have considered the effect of **reward** and **response consistency** on moral decisions. In particular, response consistency has not been investigated explicitly as it is usually a latent or observed variable in moral decision tasks.

Do people consider response consistency as part of self-consistency and moral integrity? By exploring and expanding the tasks incorporating response consistency, researchers identified the desire to behave consistently is a powerful determinant of human behavior when facing repeated situations. Consistent behavior allows one to signal one’s personal and intellectual strength^13^, thereby resulting in a strong motivation of maintaining self-image consistency. According to results from a self-consistent Bayesian model in a perceptual decision-making task, when making a decision, one’s perception is influenced by both the present sensory evidence and previous decisions^14^. Despite efforts to study consistency in perceptual decisions^15^, exploration/exploitation trade-off^15^, risk behavior^16^, the framing effect on decisions^17^, and consumer judgements^18^, the impacts of self-consistency on moral decision processes have been largely overlooked. Moreover, in moral tasks, participants were presented repeatedly with the same or similar situations^19^, but the effect of self-consistency was barely studied. These existing studies may have several unresolved questions or issues. First, most studies focus on short-term response consistency such that researchers considered the switching effect of the preceding choice as self-consistency but ignored the intertemporal consistency^14^. Further, the effect of self-consistency was only considered implicitly by the decision maker but not quantified in an explicit way^14^. That is, the cue of response history has not been presented to the participants and utilized as a factor to consider in their decisions.

At the neural level, the **reward system** is tightly linked to moral decisions. Functional magnetic resonance imaging(fMRI) studies have shown that social norm-based intrinsic rewards are encoded in the nucleus accumbens (NAcc) and caudate nucleus—the brain regions responding to external rewards such as monetary gains, thereby providing direct evidence for the existence of an intrinsic reward system for social norms^20^–^22^. Notably, the resting-state functional connectivity between the reward and moral networks can predict dishonest behavior in the later task^23^. This intrinsic reward system might affect moral behavior, when it fails to control, spontaneous dishonesty is induced^7^. **Cognitive control** also plays a crucial role in moral decision^24^ because resisting the temptation of the reward usually results in moral behaviors^7^. The “Will and the Grace” hypotheses have been at odds^25,26^. The “Will” hypothesis suggests that people are usually selfish and dishonest, and people must consciously exert cognitive control to keep moral^27^–^30^. In contrast, the “Grace” hypothesis holds that people are intuitively honest and require cognitive control to override their dominant honest impulses to cheat for self-interest and has been supported by findings that people are more honest under time pressure^31^. Speer et al.^32^ found that the “Will and Grace” hypotheses controversy can be reconciled. Cognitive control is not needed for people to be honest or dishonest per se; rather, it depends on an individual’s moral defaults (inclination to be honest or dishonest)^7^. Specifically, cheaters exhibit increased activation in the cognitive control brain regions, such as the anterior cingulate cortex (ACC) and inferior frontal gyrus (IFG), and greater activity in the nucleus accumbensnAcc when deciding to be honest. Given cognitive control’s role in congruency processing^33^, it also exerts control over consistent decisions. This research aims to gain a better understanding of cognitive control and the reward brain in (dis)honest decisions. The trade-off between consistency and reward originates from an interplay brain system, in which reward and cognitive control are essential.

Over the past decade, mouse tracking(MT) has been implemented as a powerful tool in the decision to measure real-time mouse movements^34^, which can provide information regarding the time course of a decision process, such as hesitating or changing one’s mind halfway through. It has also been applied to the detection process of dishonesty or untruthful responses^35,36^. Since stable decisions usually show the direct route in mouse trajectories, we propose that consistent decisions would show less hesitation and smoother trajectories. Regarding reaction time (RT) or MT data, the drift-diffusion model (DDM) has also been implemented to understand various binary decisions in value-based decision-making^37^–^39^. In a simple DDM, the model encodes a relative value signal that measures the accumulated “evidence” and other parameters. For example, a simple DDM is characterized by the following four parameters: (1) the symmetric location of the barriers (±b); (2) the linear slope of the drift rate dm (so that in any trial, *µ* = dm · (*v*_*left*_ -*v*_*right*_), where *v*_*left*_, *v*_*right*_ denotes the value of the items); (3) a starting position representing side bias and often fixed at zero; and (4) a latency time (Tm) that measures a fixed amount of time out for every trial prior to the initiation of the comparison process (i.e., the time that passes from the appearance of the items and the beginning of the DDM computations)^40^. Although several studies have implemented MT or DDM with fMRI to resolve the neural mechanism of certain decisions^41,42^, to our knowledge, few studies combined MT, DDM, and fMRI to quantify the latent dynamics and investigate the underlying neural mechanism of both reward-seeking and consistency-maintaining in moral decision-making.

Inspired by previous studies, we aim to investigate the role of reward and consistency in moral decision-making, which has been overlooked in previous research. With this aim, we dissect the contributions of reward and self-consistency to moral decisions by collecting evidence from behavioral (e.g., RT and MT), computational model (e.g., DDM), and brain images (e.g., fMRI) data. Specifically, we propose the following four scientific questions (Figure 1): (**Q1**) What is the role of self-interest and consistency in flexible moral decisions? (**Q2**) How are reward and consistency considerations associated with the cognitive control-related and reward-related brain regions? (**Q3**) How does decision flexibility link to mouse trajectories, and the related brain regions? (**Q4**) How are mouse trajectories and brain activity patterns during the reward and consistency trade-off mediated by individual differences? To answer these questions, we designed a binary value-based task paradigm and experimentally manipulated these variables (i.e., reward magnitude, response history, and the answer), which yielded interpretable spontaneous moral decisions. The novelty of our research lies in combining explicit response history presentation to 1) build a theoretical framework on moral decisions with considerations of self-interest, and self-consistency; 2) uncover the latent decision dynamics and quantify the evaluation between reward and consistency; and 3) dissociate the behavioral and neural representation associated with reward and consistency.

**Figure 1.**
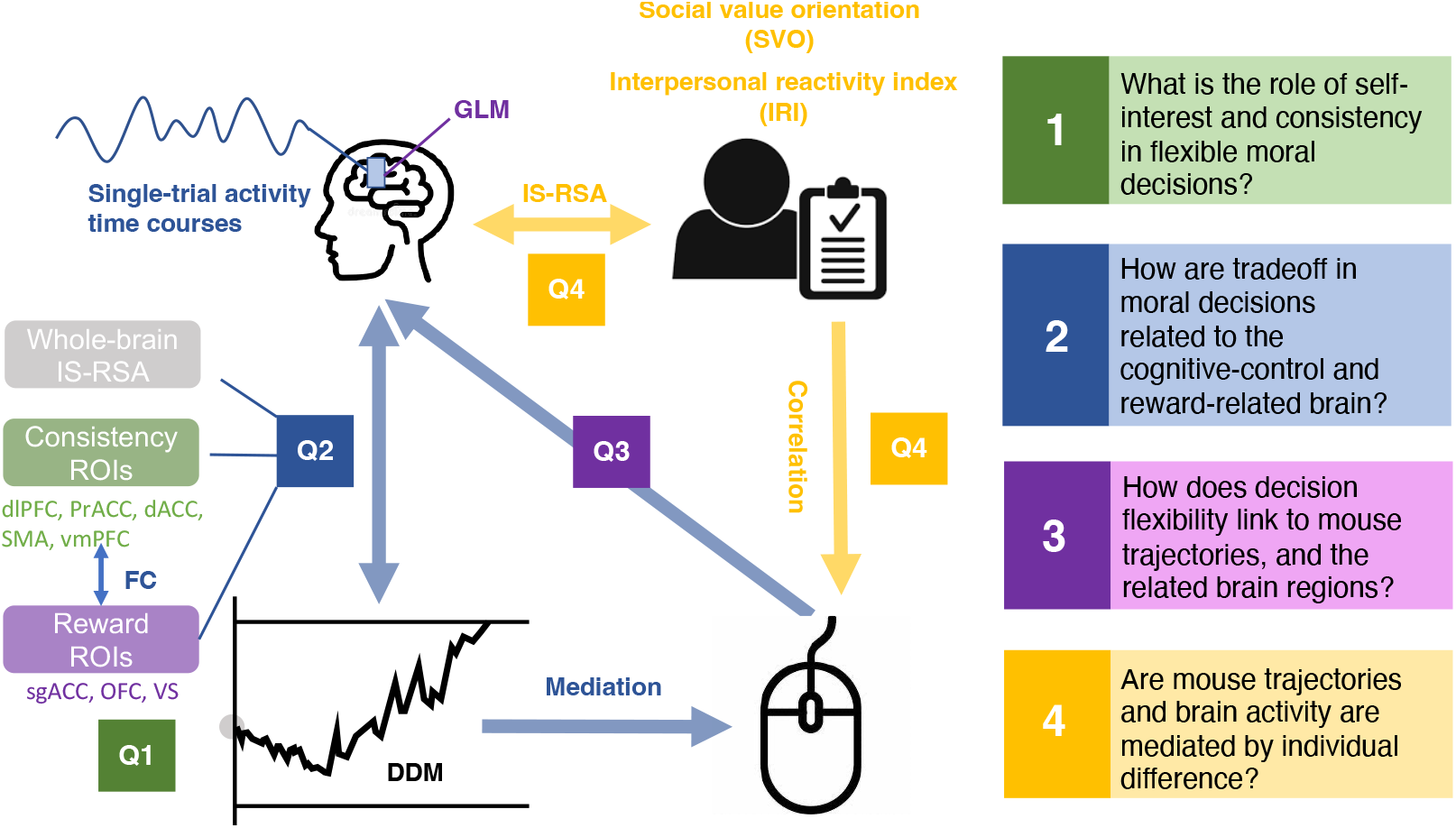
Overview of the study’s framework. This study’s framework focuses on addressing four research questions. Left: We use fMRI in combination with computational modeling and mouse tracking technique to capture the decision processes in moral decision-making. To answer **Q1, w**e fit a tDDM to behavioral data to estimate parameters (i.e., weights and latencies of consistency and reward), participant by participant. To answer **Q2**, we correlated the tDDM parameters with univariate activity, neural RDM, and functional connectivity within priori ROIs. To answer **Q3**, we defined decision entropy of every participant to quantify flexibility and correlated it with mouse tracking indices and univariate activity. Finally, to answer **Q4**, we performed IS-RSA to identify the individual variations in brain activity. Also, we delved into the relationship between neural activity within the ROIs and mouse tracking indices and participants’ personal traits. Right: four research questions of the study.

## Results

### Behavioral Results

The participants ^1^ completed the self-paced information transmission task (see *Methods* section for details) inside the MRI scanner. They were asked to delivery information to another person who was playing a correct-answer game. The task comprised nine runs with 20 repeated uncommon questions (e.g., When do penguins usually lay their eggs?) presented in the middle of the screen, with the correct and erroneous answers presented in the upper corners (see Figure 2a). Simultaneously, corresponding monetary rewards (a red number) and the previous choices (blue triangles) were also presented under the answers. The participants were instructed to decide which answers to deliver while the other person had no information about the correct answer; thus, they were more likely to trust the delivered answer. The dishonesty probability was computed as the ratio of selecting wrong answers. The behavioral results showed large individual differences in the total levels of dishonesty (mean = 41.12%, median = 43.21%, SD = 18.23%; Figure 2b).

**Figure 2.**
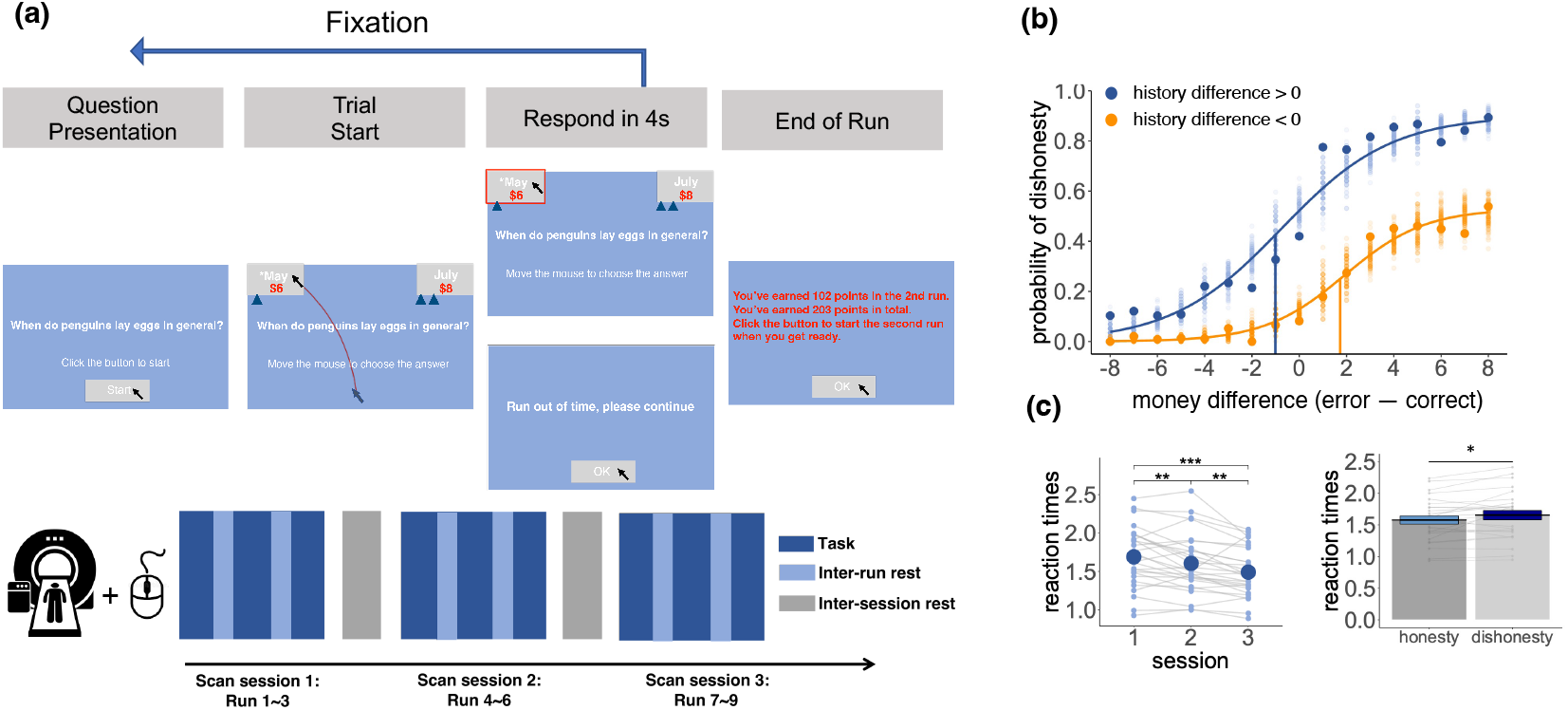
Illustration of the experimental paradigm and behavioral results. **(a)** The task procedure . The participants were presented with trivial questions, after which they were presented with two answers in the top left or top right corner, respectively, with different amounts of reward. They were asked to deliver one answer to another person with the goal of gaining more rewards. For the example trial, the left answer (with *) was correct but assigned a lower reward coin (’6’), whereas the other answer was incorrect, but assigned a higher reward (’8’). The entire task comprised three scanning sessions, each including three runs. **(b)** The dishonesty probability changes as a function of response history and reward difference. We fitted the psychometric curve along the reward difference (error minus correct choice) under two conditions at the group level: One was when participants’ history of dishonest choices was more than that of their honest choices and the other condition was the opposite. We found that history of choices not only shifted the threshold but also the total dishonesty rate of the psychometric curve across different reward difference trials. **(c)** How reaction times were affected by the session and responses. We observed the linear effect of task sessions. Participants took longer RTs when making dishonest choices.

First, we explored how choice history contributed to participants’ current choices. We fitted the psychometric curve under two conditions on the group level. History difference referred to the number of dishonesty choices minus the number of honesty choices for a certain question. A positive history difference indicated that participants lied more when faced with a certain question. We found that the history of choices shifted both the threshold and the total dishonesty rate of the psychometric curve (Figure 2b).

Next, we explored whether reaction times (RTs) can be predicted by the session and responses. Consistent with the Grace hypothesis that dishonesty requires cognitive processing, the participants spent more time making dishonest choices (paired t-test: *t*_(26)_ = 2.50; *p* = 0.02; 95% confidence interval (CI) = -0.14 to -0.01). Additionally, we estimated a mixed-effect linear regression model using the participants’ RT as the dependent variable and the session as the independent variable. We observed that reaction times decreased along the sessions as indicated by a negative task session effect (*t*_(27,4531)_ = 4.27; *p <* 0.001; regression coefficient *β* = 0.10; 95% CI = -0.15 to -0.05; Figure 2c).

#### Mouse tracking patterns along the sessions

We first plotted the trajectories under each condition (dishonesty/honesty) for three sessions (Figure 3a).The preprocessed trajectories were remapped to the left side and temporally normalized to 101 time points. We split the trajectories into dishonesty and honesty conditions for each session (Figure 3a), and computed paired t tests between the two conditions on each time point. The results showed that in the last session, the trajectories under both conditions began to present a large window of significant differentiation (time points from 33 to 89, *p <* 0.01 FDR corrected). In the first session, the differentiation only spanned from the 54rd to the 62th time points (*p <* 0.01 FDR corrected, see the orange windows in Figure 3a).

**Figure 3.**
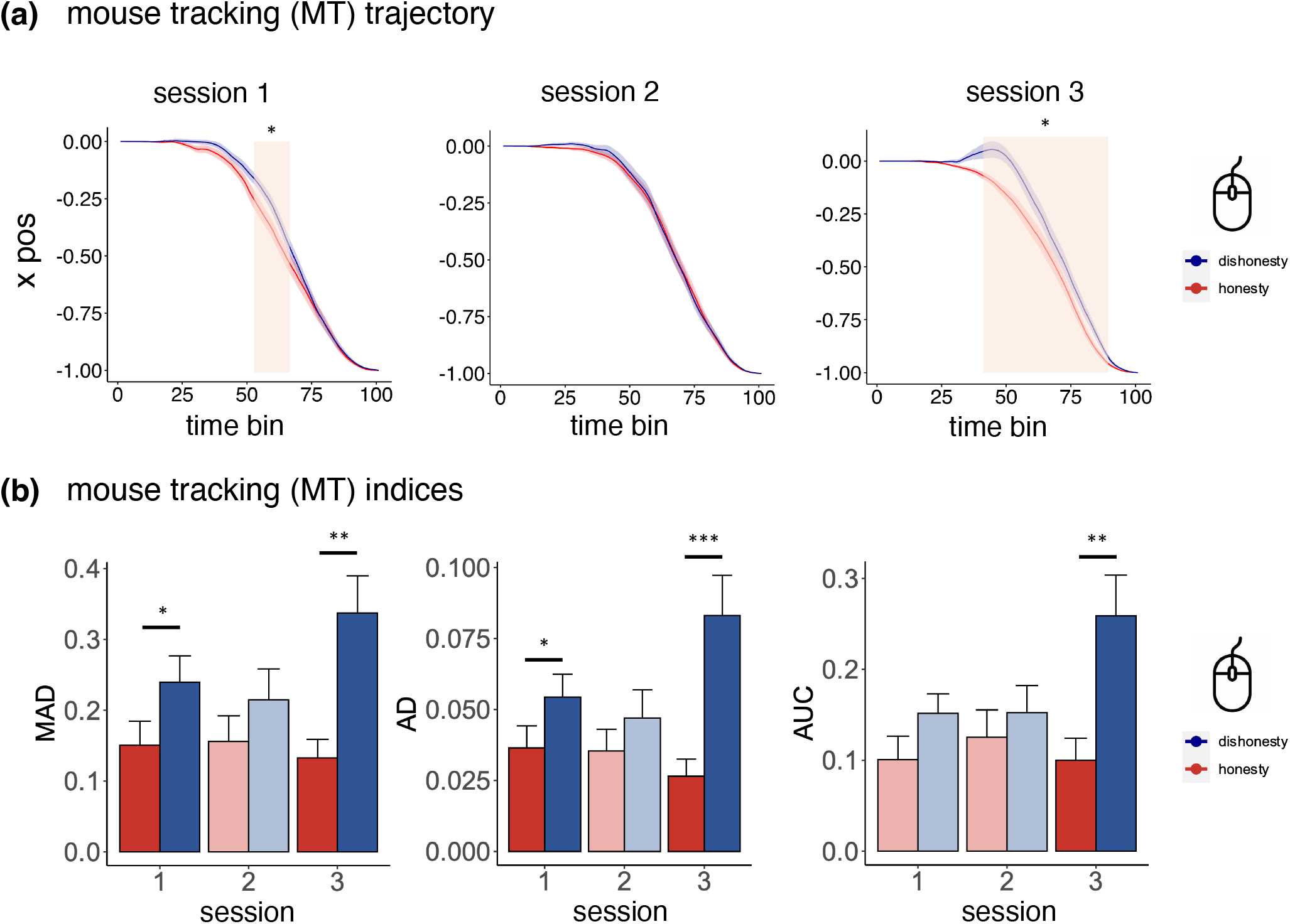
Investigations of mouse tracking. **(a)** The averaged mouse trajectories of x positions for honesty (blue) and dishonesty (red) trials. Orange windows indicate the time points that dishonesty and honesty condition exhibits significant different x positions. **(b)** Under both session 1 and session 3, dishonesty trials has significantly larger trajectory deviation than honesty trials (* indicates *p <* 0.05, ** indicates *p <* 0.01,*** indicates *p <* 0.001). Error bars indicate between-subject standard error.

Several different measures (maximum absolute deviation (MAD), average deviation from the direct path (AD), and area under the curve (AUC)) for the curvature of mouse trajectories were calculated to investigate whether any static index could reflect differences between the dishonest and honest responses. A paired t-test was conducted on these measures under different conditions in the three sessions, and the results showed in the last session that these indices presented a more significant difference between the dishonesty and honesty conditions (MAD in session 1: *t*_(26)_ = 2.33, *p* = 0.03, 95% CI = 0.01 to 0.17; MAD in session 3: *t*_(26)_ = 3.67, *p* = 0.001, 95% CI = 0.09 to 0.32; AD in session 1: *t*_(26)_ = 2.17, *p* = 0.04, 95% CI = 0.000943 to 0.035; AD in session 3: *t*_(26)_ = 3.85, *p <* 0.001, 95% CI = 0.026 to 0.087; AUC in session 3: *t*_(26)_ = 3.38, *p* = 0.002, 95% CI = 0.06 to 0.26) (see Figure 3b). As an index of the conflict levels^43^, the AUC results may indicate making a dishonest decision considering that both choice history and reward are more difficult than making an honest response.

#### DDM results: Reward and consistency had biased contributions and latencies in decisions

Previous studies^44,45^ found a dissociated influence of different factors on decision-making using time-varying drift-diffusion models. Similarly, we constructed four different DDM models (see *Methods* for details) with different drift rates and/or latencies. We identified that the tDDM, in which reward and consistency had different drift rates and started influencing the decision-making process at different onset times) performed best over three other models. The tDDM showed a significantly lower Bayesian information criterion (BIC) than the three alternative models (tDDM-mDDM: *t*_(26)_ = -10.66, *p <* 0.001, 95% CI = -272.77 to -184.60; tDDM-sDDM: *t*_(26)_ = -12.20, *p <* 0.001, 95% CI = 345.51 to -245.85; tDDM-latDDM: *t*_(26)_ = -4.83, *p <* 0.001, 95% CI = -91.96 to -37.02; for detailed statistical information of DDMs, see Table **??** in the *Supplementary materials*). As the latDDM had second-lowest mean BIC score, we compared the tDDM with the latDDM for each participant. Critically, although BIC penalizes tDDM for having one more latency parameter, the tDDM performed better for nearly all participants (26 out of 27 participants; Figure 4c). Additionally, there was no statistically significant correlation between reward and consistency drift rates (Pearson *r* = −l 0.12, *p* = 0.54), nor between reward and consistency latencies (Pearson *r* = 0.36, *p* = 0.066. Each attribute’s drift rate and latency were not statistically significantly correlated (**??** in the *Supplementary materials*). The results validated the necessity of including independent drift rate and latency parameters for reward and consistency.

**Figure 4.**
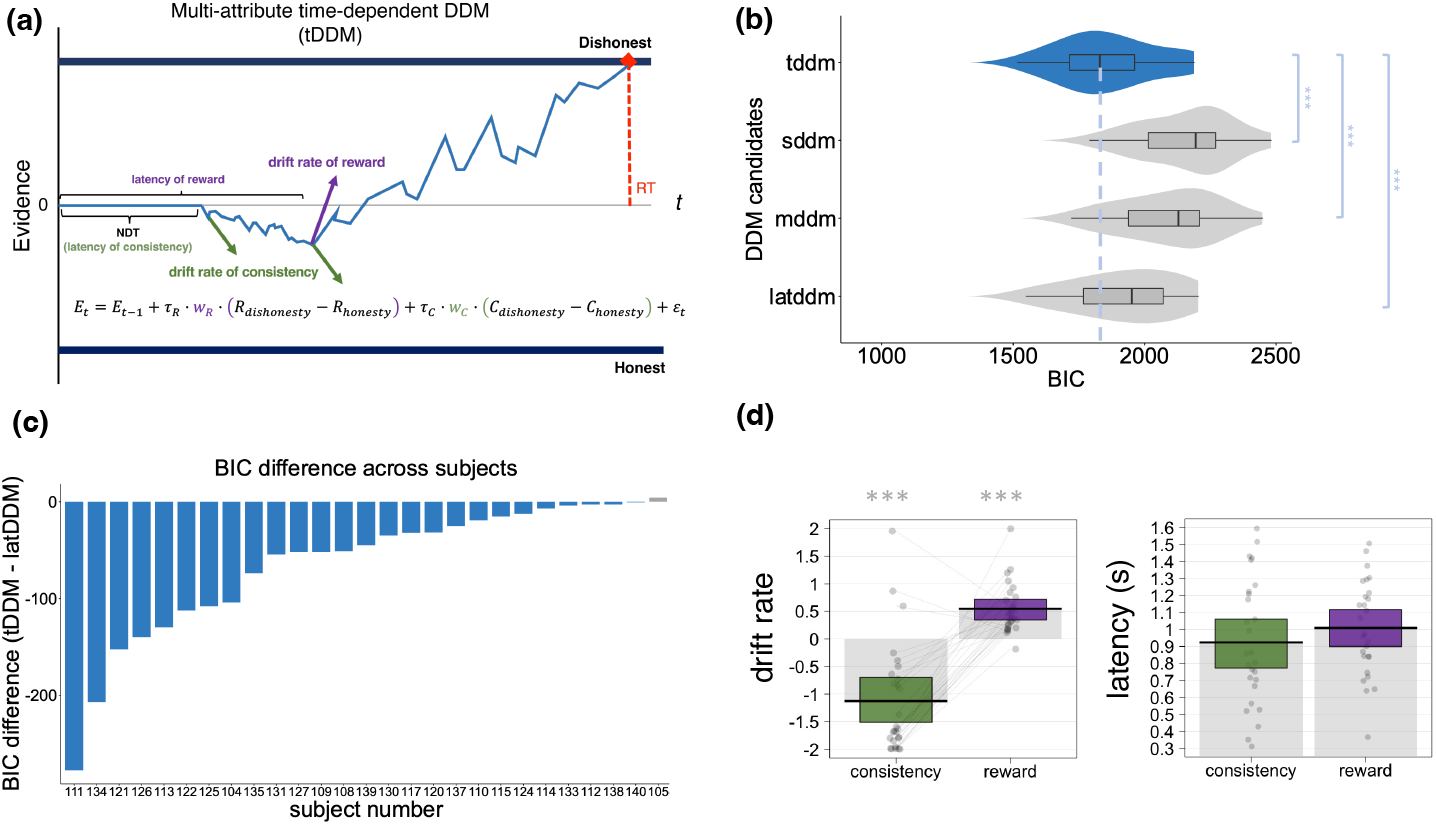
The construction of multi-attributes time-dependent DDM (tDDM), model comparison and fitting results. **(a)** The illustration of the tDDM. The tDDM assumed reward and consistency (two considered attributes in decision making) and started to affect the evidence accumulation process with different drift rates and at different onset timings (reward: purple; consistency: green). **(b)** Model comparison results. The tDDM was the winning model, with a BIC value that was significantly smaller than that of the other alternative DDMs. The significance stars show the p-values of the difference effects (*** *p <* .001). **(c)** Summary of the individual-level model comparison results of tDDM and latDDM (only worse than tDDM). Nearly all the participants showed better fitting with tDDM. **(d)** Fitted parameters for consistency and reward. The drift rate for reward was significantly higher than the drift rate of consistency (left), which drove the participants to make dishonest choices. Latency parameters between reward and consistency did not show statistical significance, but consistency came to affect the evidence accumulation process earlier than reward.

We then tested the significance of each parameter with the tDDM fitting results. We found that both reward and consistency significantly impacted the decision-making process (reward drift rate: *t*_(26)_ = 6.33, *p <* 0.001, 95% CI = 0.37 to 0.73; consistency drift rate: *t*_(26)_ = -5.8, *p <* 0.001, 95% CI = -1.53 to -0.73; Figure 4d)). However, the consistency drift rate was negative (mean = -1.13, sd = 1.01; Figure 4d, left panel)), indicating that maintaining consistency was more likely to drive participants to be honest, which confirmed the Grace hypothesis that people were intuitively honest. The latencies of reward and consistency did not differ significantly. However, they showed a large variation (reward: mean = 1.01, sd = 0.28; consistency: mean = 0.92, sd = 0.37; Figure 4d, right panel), thereby indicating that consistency and reward had different durations of impact on the decision-making process. Together, these results shed light on how the participants evaluated monetary reward and choice history. None of the two attributes had an absolute advantage over the other, thereby confirming the competing relationship between the two.

To validate the model, parameter recovery tests were performed to ensure that the competing advantage did not lead to inaccurate decisions compared to the empirical data. Choice and RT patterns were simulated using the best fitting parameters of the tDDM based on the consistency and reward differences participants faced in each decision trial. The simulated choices reached over 81.62% accuracy according to the empirical data, and the simulated RTs under dishonesty and honesty conditions were close to the real RTs (real RT minus simulated RT in dishonesty condition: mean = 0.41, 95% HDI= [-2.97, 4.34]; real RT minus simulated RT in the honesty condition: mean = -0.96, 95% HDI= [-5.90, 1.73]; Figure **??**c in *Supplementary materials*).

The same fitting procedure was applied to the generated data. Consistent with the results of fitting the empirical data, the tDDM performed best among the four DDMs, as assessed using the lower Bayesian information criterion (BIC; Figure **??**a in *Supplementary materials*). Apart from the tDDM, the mDDM performed best. We thus compared the BIC of the tDDM with the mDDM of each participant and found that the tDDM performed better for every single participant (Figure **??**b in *Supplementary materials*). Furthermore, our recovery procedure for the tDDM yielded accurate parameter estimates. Critically, the parameter recovery tests also indicated a significantly higher (and positive) drift rate for reward and negative drift rate for consistency. Different latencies were also identified (Figure **??**c in *Supplementary materials*). Correlation was conducted between the recovered and original parameters. Significantly high correlations were found for all four parameters (consistency drift rate: *r* = 0.98, *p <* 0.001; reward drift rate: *r* = 0.99, *p <* 0.001; latency of consistency: *r* = 0.86, *p <* 0.001; latency of reward: *r* = 0.93, *p <* 0.001; Figure **??**a and b in *Supplementary materials*).

### Univariate analysis results

To characterize the neural processes underlying the competing relationship between consistency and reward in moral decision-making, we attempted to dissect the two processes and their links with the fMRI data. According to previous studies^31,32^,46, dishonesty required cognitive control to override their dominant honest impulses to cheat for profit. We mainly focused on the cognitive control- and reward-related brain regions. To simply identify whether our paradigm triggered activation in specific regions, we estimated a whole-brain GLM, which included *response type* (dishonest / honest) for each session, with the RTs as durations and modulators^47^. To better probe the trading-off process between consistency and reward, the last session was selected for analysis. Dishonesty responses identified regions associated with cognitive-control (i.e., dlPFC, IFG, SMA and prACC) and reward (i.e., putamen) as expected (Figure **??** in *Supplementary materials*; for detailed activation table of dishonesty responses in session 3, see Table S**??** in *Supplementary materials*). This indicated that our experiment setting successfully induced the intention to lie for self-reward. Next, we investigated whether the activation in cognitive control related regions was unique in the last session because it required a full trade-off between reward and consistency as history choices had mounted to the maximu. We selected several ROIs (i.e., cognitive control brain regions, reward brain etc.) with masks (Figure 5a) of these regions from Neurosynth’s (https://neurosynth.org/) meta-analysis and Brodmann atlas^48^. Beta maps of the first-level GLM within the masks were extracted for each condition on the subject level. In SMA, PCC, and dACC, we found that the activity under the condition of honesty was stronger in the last session compared to the first session (SMA: *t*_(26)_ = 3.53, *p* = 0.002, 95% CI = 0.13 to 0.503; PCC: *t*_(26)_ = 2.16, *p* = 0.04, 95% CI = 0.01 to 0.48; dACC: *t*_(26)_ = 3.12, *p* = 0.005, 95% CI = 0.11 to 0.53; Figure 5b). For dlPFC and prACC, both dishonesty and honesty conditions showed increased, though non-significant, activity (dlPFC in dishonesty condition: *t*_(26)_ = 1.59, *p* = 0.12, 95% CI = -0.06 to 0.48; dlPFC in honesty condition: *t*_(26)_ = 1.045, *p* = 0.31, 95% CI = -0.16 to 0.495; prACC in dishonesty condition: *t*_(26)_ = 2.03, *p* = 0.05, 95% CI = − 0.0025 to 0.4005; prACC in honesty condition: *t*_(26)_ = 1.50, *p* = 0.15, 95% CI = -0.08 to 0.54; see Figure 5b).

**Figure 5.**
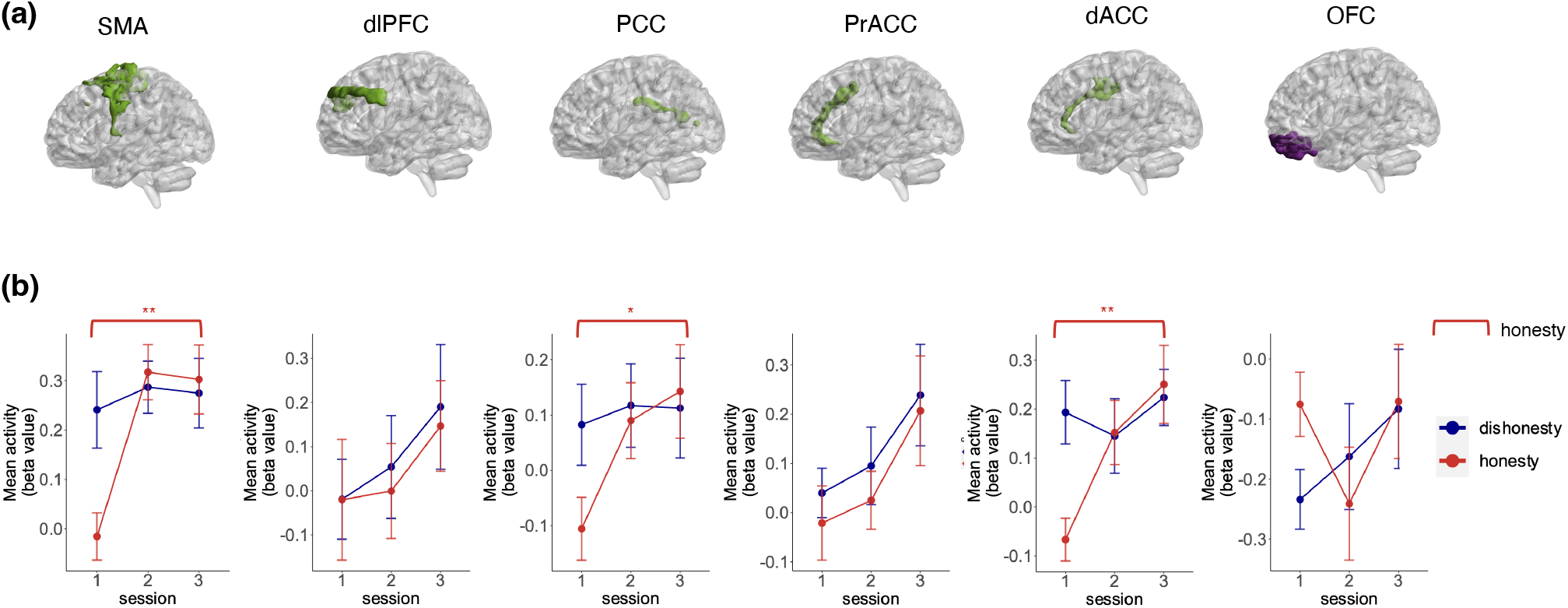
Univariate results in session 3. **(a)** The ROIs for consistency (green) and reward (purple). Masks of these ROIs are obtained from Brodmann atlas^48^ and Neurosynth (https://neurosynth.org/). **(b)** Changes of activations along the sessions in regions of interest. The activity of dlPFC and prACC increased as the session progressed for both conditions. No significant changes in activation were observed in dlPFC or prACC across sessions (* indicates *p <* 0.05, ** indicates *p <* 0.01). Error bars indicate between-subject standard error.

### Drift rates of consistency and reward modulated the activity of cognitive-control and reward regions

To explore the neural basis of the evaluation of consistency and reward, we extracted the mean betas from the pre-defined ROIs and performed correlation analysis with model drift rates (Figure 1b). The activity in SMA in session 3 was correlated with the drift rates of both consistency and reward (correlation with the drift rate of consistency: dishonesty: Pearson’s *r* = -0.51, *p* = 0.007, 95% CI =-0.75 to -0.16; honesty: Pearson’s *r* = -0.39, *p* = 0.04, 95% CI = -0.67 to -0.01; correlation with the drift rate of reward: dishonesty: Pearson’s *r* = 0.37, *p* = 0.06, 95% CI = -0.02 to 0.65; honesty: Pearson’s *r* = 0.41, *p* = 0.03, 95% CI = 0.04 to 0.68; see Figure 6a, left and middle panel). For other consistency ROIs like vmPFC, dlPFC, and prACC, the higher the weight on consistency, the higher the activity in these regions under at least one condition (vmPFC under dishonesty condition: Pearson’s *r* = 0.41, *p* = 0.03, 95% CI = 0.04 to 0.68; vmPFC under honesty condition: Pearson’s *r* = 0.44, *p* = 0.02, 95% CI = 0.07 to 0.702; dlPFC under honesty condition: Pearson’s *r* = 0.41, *p* = 0.03, 95% CI = 0.04 to 0.69; prACC under honesty condition: Pearson’s *r* = 0.43, *p* = 0.03, 95% CI = 0.06 to 0.697; see Figure 6b first and the third columns). For dACC, drift rate of consistency was negatively correlated with its contrast activity (dishonesty *>* honesty) (Pearson’s *r* = -0.65, *p <* 0.001, 95% CI = -0.82 to -0.35; see Figure 6b right lower panel).

**Figure 6.**
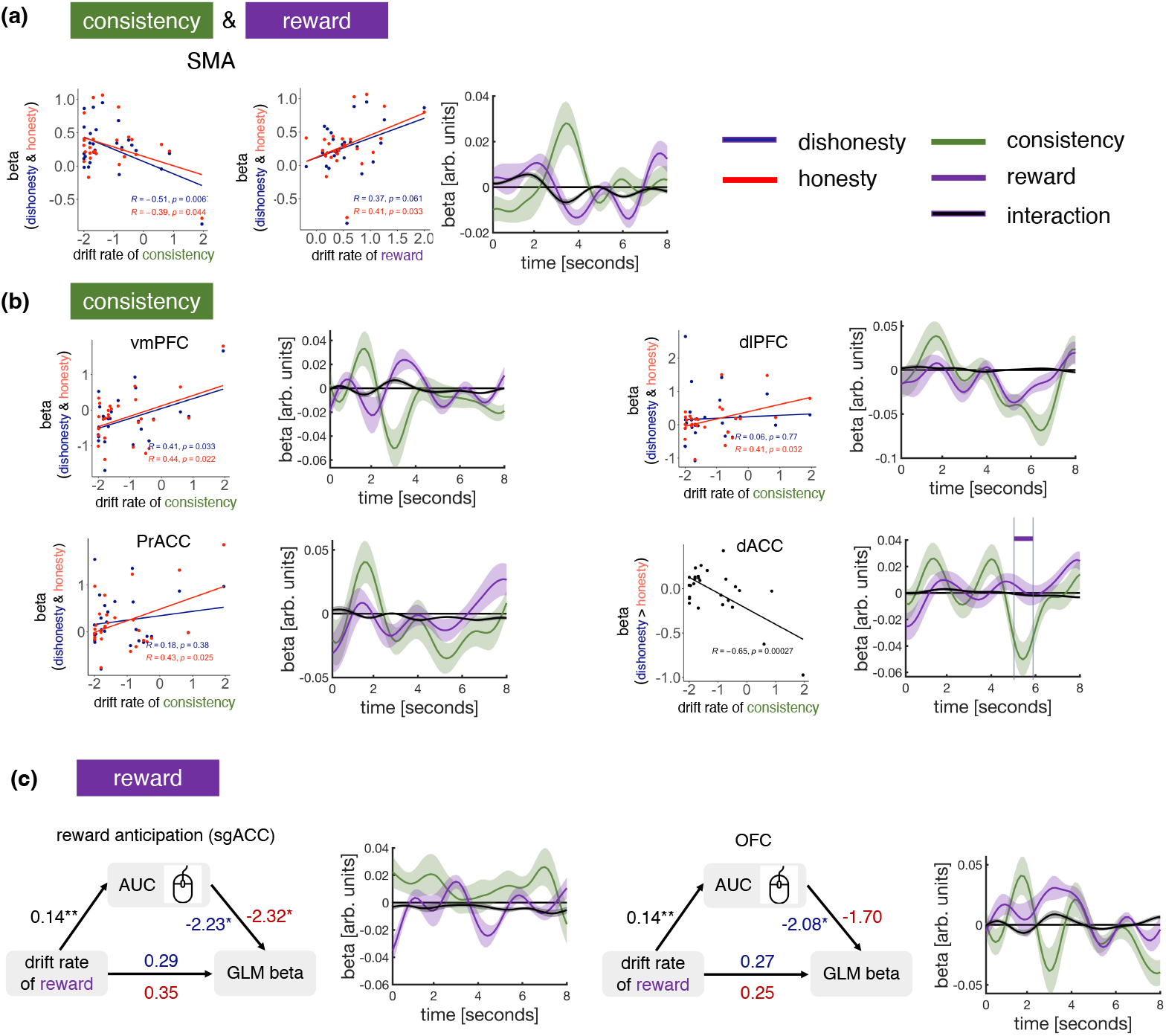
Relationships between tDDM parameters and ROI activity. **(a)** Brain activity within SMA in session 3 showed correlations with both drift rate of consistency and reward. Red color represents the honesty condition, and blue color represents the dishonesty condition. Last figure: The time courses are coefficients from a regression in which we predicted SMA activity time courses using reward difference (purple), consistency difference (green) and the interaction of these two (black). **(b)** Brain activity associated with consistency. Activation in vmPFC in session 3 increased with the drift rate of consistency for both conditions. There was a positive correlation between the drift rate of consistency and activation in dlPFC or prACC in session 3 only for honesty condition. There was no significant correlation in relationship between drift rate of consistency and activation in dlPFC or prACC in session 3 for the dishonesty condition. Second and last columns: We predicted the single-trial time courses of vmPFC in session 3 using reward difference (purple), consistency difference (green) and the interaction of the two (black). We predicted the time courses of dlPFC, prACC, and dACC using the interaction of response (dishonesty = 1, honesty = 0) and consistency difference (green), the interaction of response and reward difference (purple) and the interaction of reward and consistency difference (black). **(c)** The AUC index of MT mediates the relationship between drift rate of reward and brain activity. First and third columns: The MT index AUC fully mediated the relationship between drift rate of reward and brain activity of reward-related ROIs (* indicates *p <* 0.05, ** indicates *p <* 0.01). Second and last columns: The time courses were predicted in the same procedure as SMA.

However, the reward drift rate was not directly associated with the reward ROIs. Their relationships were fully mediated by the mouse tracking index AUC (Figure 6c). AUC represents the geometric area from the reference trajectory to the actual trajectory. The higher the AUC, the bigger the attraction to the other response option. AUC is a good indicator of the overall attraction^49^; thus, we analyzed two reward-related ROIs. One was the reward anticipation mask obtained from Neurosynth (z ¿10); the other was the OFC derived from the Brodmann atlas^48^. For anticipation ROI, AUC mediated the relationship between the reward drift rate and the GLM mean betas under both honesty and dishonesty conditions. Regarding OFC, AUC played a mediating role only in the dishonesty condition.

The results identified that SMA was the only ROI correlated with both attributes (consistency and reward). SMA was negatively correlated with the consistency drift rate in both responses (honesty: Pearson’s *r* = 0.39, *p* = 0.04, 95% CI = -0.67 to -0.01; dishonesty: Pearson’s *r* = -0.51, *p* = 0.007, 95% CI = -0.75 to -0.16) and positively correlated with the drift rate of reward in honesty responses (honesty: Pearson’s *r* = 0.41, *p* = 0.03, 95% CI = 0.038 to 0.68; dishonesty: Pearson’s *r* = 0.37, *p* = 0.06, 95% CI = -0.02 to 0.65; left two figures in Figure 6a).

It is notable that the tDDM not only estimated the weights on consistency and reward but also offered timing of when the two attributes affected the decision making. Therefore, to gain further insight into the dynamics of brain activity association, we also quantified the neural impact of consistency and reward in the time domain. Specifically, we extracted trial-wise activity estimates (from session 3) in an 8s window from the onset times within the ROI masks. Then, we regressed the preprocessed time courses on each time point over several regressors, including consistency difference, reward difference, and their interaction. The results showed that SMA tracked consistency, reward, and their interaction but tracked reward and the interaction first (Figure 6a third column). For cognitive-control-related ROIs, they mainly tracked the consistency drift rate (Figure 6b, second and fourth columns). Specifically, vmPFC also encoded reward but later than it did consistency, which was in line with t he previous research showing that vmPFC encoded value ^50^–^52^ For reward-related regions, reward anticipation ROI encoded both reward and consistency, while OFC exhibited a more stable encoding of reward (Figure 6c, second and fourth columns).

### Individual variations in brain activity reflect differences in evaluation between reward and consistency

We observed heterogeneity in participants’ dishonesty rates, with some individuals lying more frequently, while others seldom lied (Figure **??**). Moreover, participants’ weights on consistency and reward formed a two-dimensional space (Figure 7a, right panel), in which the dots were distributed across at least two quadrants and showed no correlation (see *supplementary materials Table* **??**). We suggested that different participants exhibited a variety of brain activity in the evaluation of consistency and reward. Thus, we further used inter-subject representation similarity analysis (IS-RSA) to seek the brain regions encoding the evaluation of both consistency and reward. We conducted the same analysis for dishonesty and honesty conditions separately.

**Figure 7.**
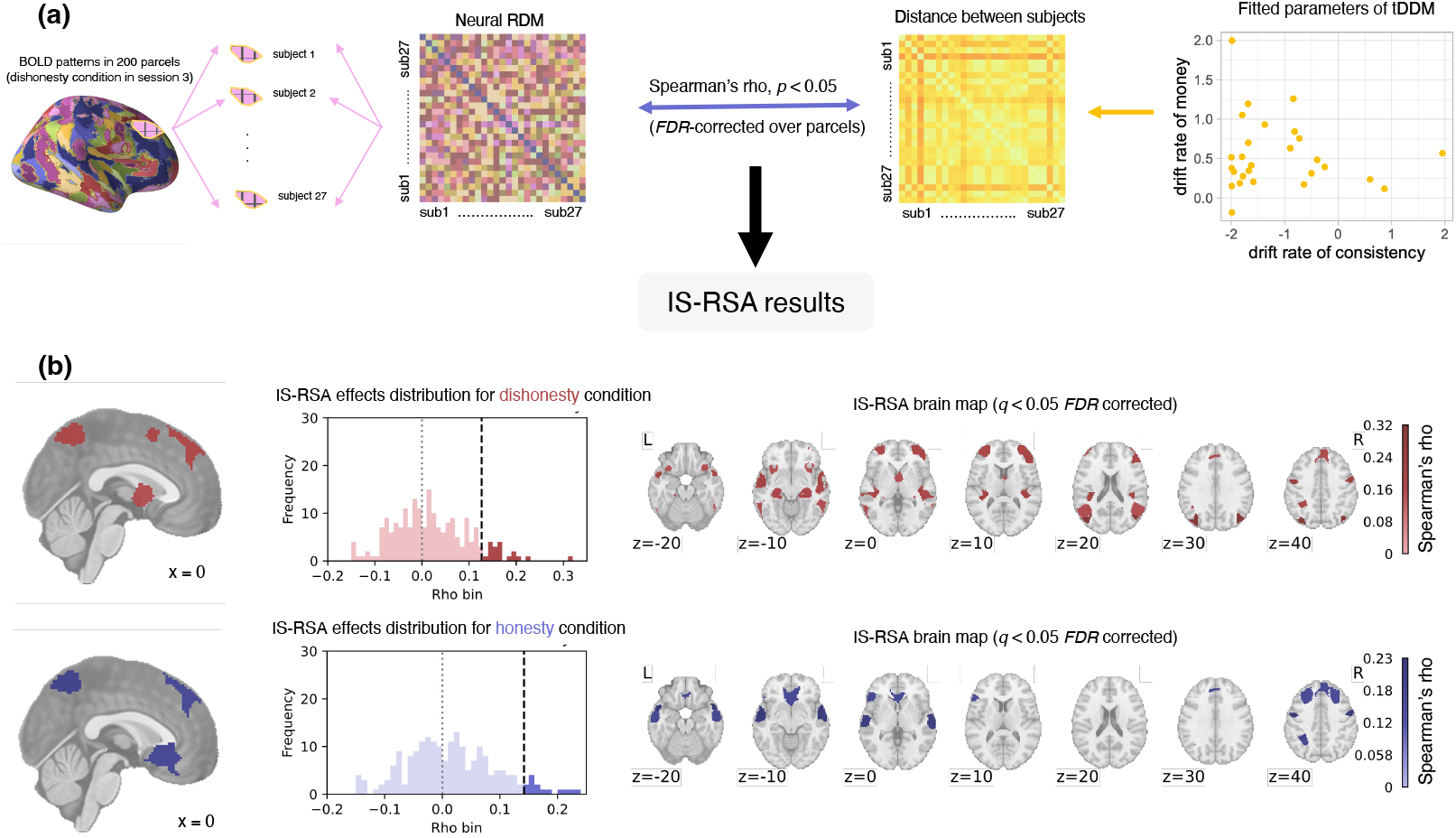
Illustration of the inter-subject representational similarity analysis and the results. **(a)** Illustration of the IS-RSA. We created a neural representational dissimilarity matrix (RDM) by calculating the distance (1-correlation) of brain activity patterns across participants for each of the 200 parcels. The behavioral RDM in the right side (yellow) was calculated by the pairwise Euclidean distance between drift rates of consistency and reward in the two-dimensional space. **(b)** The IS-RSA results. We identified parcels that survived FDR correction under dishonesty and honesty conditions. The scale represents Spearman’s rho. Overall, we observed significant brain regions over SMA, prACC and IFG for both conditions.

We first created a geometric representational space of tDDM parameter space (drift rates of consistency and reward; Figure 7a, right panel). To search for brain regions that showed a similar representational geometry to this distance measure, we used a priori 200-parcel whole-brain parcellation^53^, and calculated multi-voxel activity pattern (beta maps derived from the GLM) distances (1-correlation) between each pair of participants during session 3 for dishonesty and honesty responses, respectively (Figure 7a, left panel). This yielded one representational dissimilarity matrix for each condition. After correlating the behavioral distance matrix with the brain dissimilarity matrix, we identified parcels that survived FDR correction (*p <* 0.05). We observed significant inter-subject representational similarity effects in 18 brain parcels under the dishonesty condition, including SMA, dlPFC, mPFC, ANG, IFG, and OTC (Figure 7b, upper panel). Moreover, 14 parcels were observed under the honesty condition, including pre-SMA, IFG, prACC, SMA, MTG, and STG (Figure 7b, lower panel). These results indicated that decision-related activity patterns in these regions were more similar than in other regions across participants who shared similar weights after making repeated dishonesty choices.

### Functional connectivity explains the drift rates and personal traits

We further tested whether there was task-related functional connectivity (FC) between the cognitive control and reward brain regions that could reflect the variation of the drift rates and personal traits. Previous studies have demonstrated that dlPFC is closely related to cognitive control, and the brain lesion^54^–^56^ in dlPFC can increase dishonesty, while stimulation over dlPFC can decrease it^57^. Previous studies revealed that dlPFC interacts with the reward-related ventral striatum (VS) and conflict-related anterior cingulate cortex (ACC) to control human behavior in different situations^58^. We then tested how the FCs between dlPFC and prACC, dlPFC, and VS (Figure 8b) were modulated by behavioral traits, such as drift rates and questionnaire scores (IRI and SVO; see *Methods* for details).

**Figure 8.**
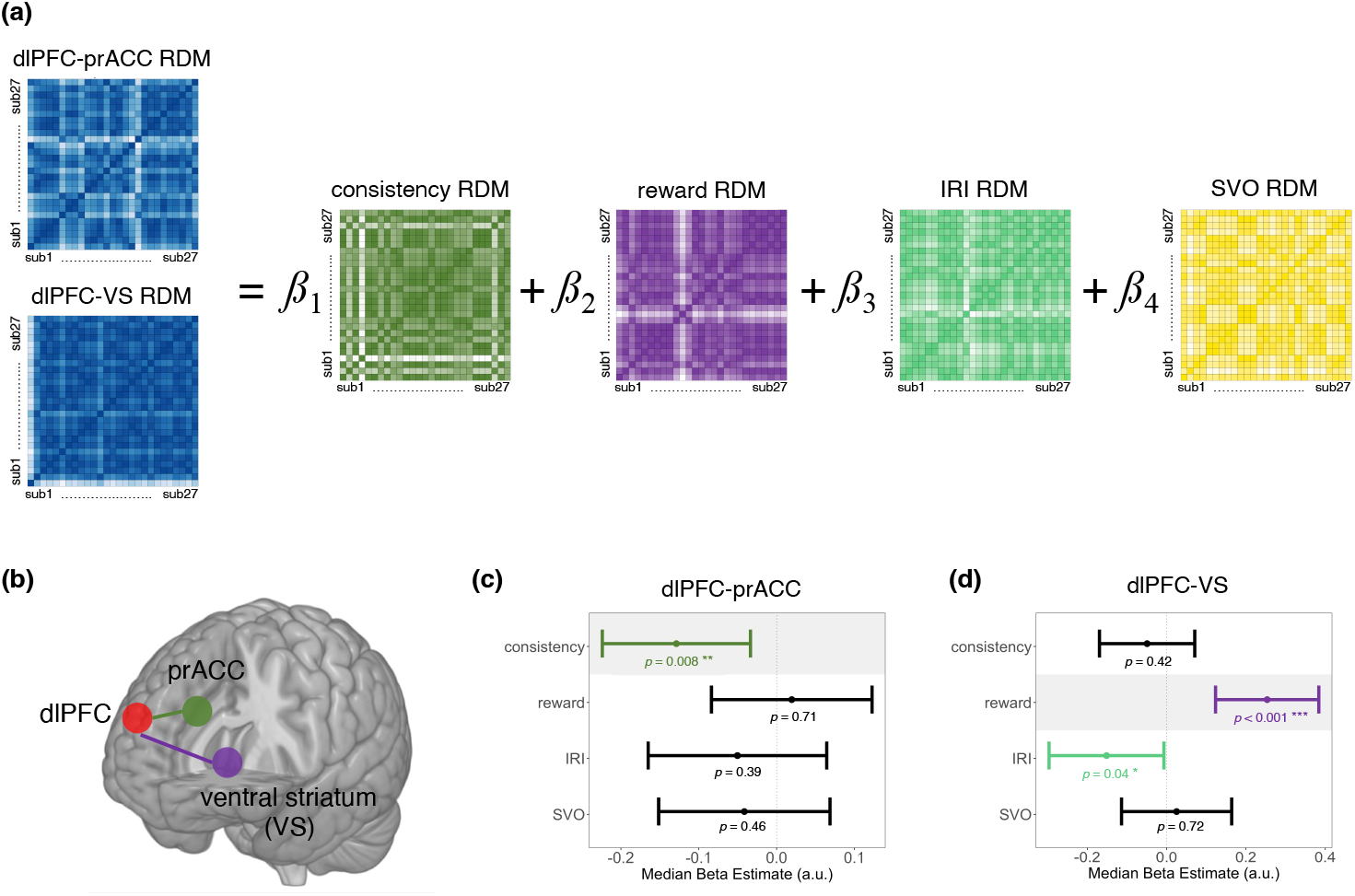
The functional connectivity analysis and results. **(a)** The regression model to predict RDM with FC of dlPFC-prACC and dlPFC-VS. We also created four predictor matrices reflecting the participants’ drift rate of consistency in tDDM (dark green), the participants’ drift rate of reward in tDDM (purple), the participants’ scores in IRI (light green), and the RDM of the participants’ scores in SVO (yellow). We built a prediction model to test how those behavioral traits shaped their variations in these two FCs (dlPFC-prACC, and dlPFC-VS), respectively. **(b)** The ROIs (dlPFC, prACC, and VS) for FC are shown for illustration. **(c)** and **(d)** The representative results for linear regression. In all the plots, the data points reflected median estimates, and error bars reflected uncorrected nonparametric 95% CIs. **(c)** For dlPFC-prACC, the similarity of the drift rates of consistency could significantly predict the similarity of FCs among the participants. **(d)** As for the dlPFC-VS, both the drift rates of reward (purple line) and the IRI scores (green line) could significantly predict the similarity of the FC (* indicates *p <* 0.05, ** indicates *p <* 0.01, *** indicates *p <* 0.001).

To better depict the relationship between FCs and behavioral traits, we implemented a linear regression model to explore the extent to which behavioral traits could predict FC variation. We extracted the two groups of FCs mentioned above for each participant using the CONN toolbox^59^ and calculated the interpersonal Euclidean distance. We also created four-predictor behavioral RDMs by using the drift rate of consistency, the drift rate of reward, IRI scores, and SVO scores, respectively (Figure 9a). Because there were different scales, we computed the predictor matrices using standardized Euclidean distance (see *Methods*). The results showed that the consistency drift rate predicted the dlPFC-prACC connectivity among the four predictor matrices with the biggest absolute coefficient value (*β* = -0.13, *p* = 0.008, 95% CI = -0.22 to -0.03; Figure 8c, left panel). For dlPFC-VS, as expected, the RDM of the reward drift rate and the IRI score significantly predicted the reward RDM (reward: *β* = 0.25, *p <* 0.001, 95% CI = 0.12 to 0.38; IRI: *β* = -0.15, *p* = 0.04, 95% CI = -0.296 to -0.0069; Figure 8c, left panel of). These results also further confirmed the validation of the tDDM.

**Figure 9.**
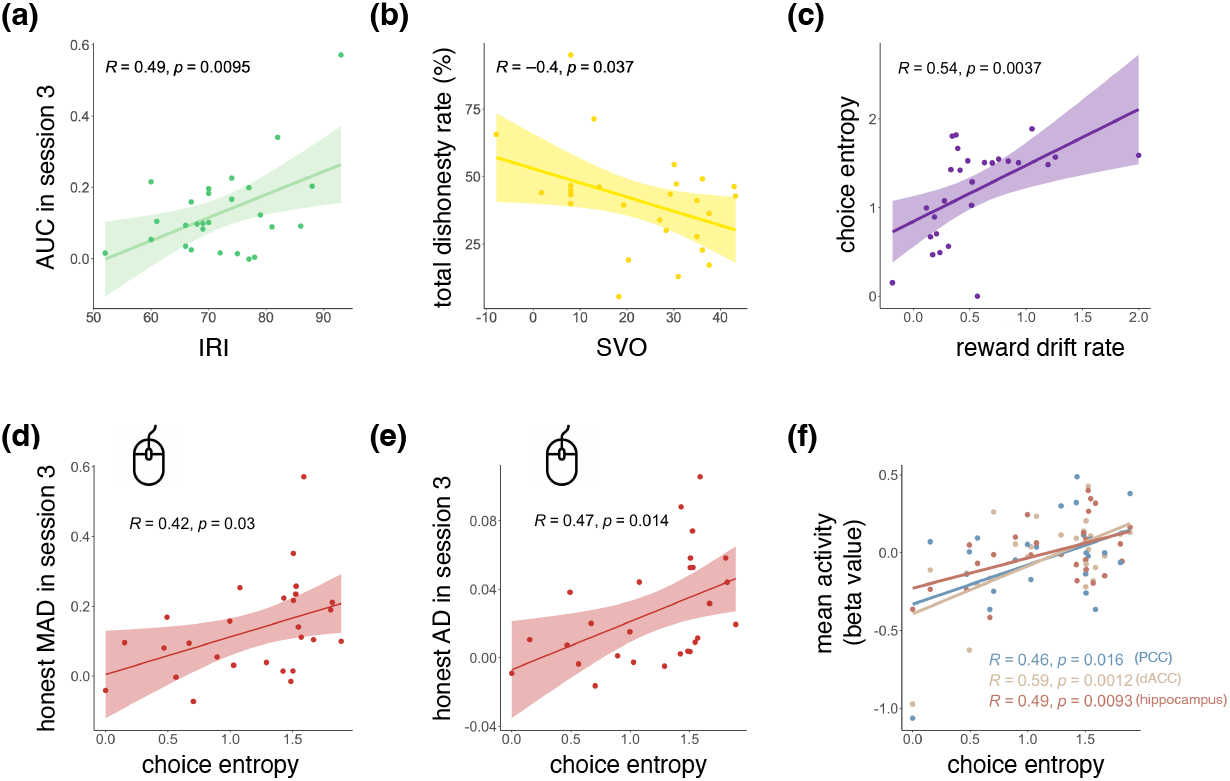
**(a)** Mean AUC of session 3 was positively correlated with interpersonal reactivity index (IRI) score. **(b)** The social value orientation (SVO) score was negatively correlated with the total dishonesty rate. **(c)** The choice entropy was positively correlated with the reward drift rate. **(c)** and **(d)** The choice entropy was positively correlated with the mouse tracking indices MAD and AD. **(e)** Mean activity (dishonesty -honesty) of PCC, dACC and hipppocampus in session 3 were positively correlated with the overall choice entropy.

### Mouse tracking and brain activity patterns were related to personal traits

We further investigated whether the MT measures and other behavioral results were associated with personal traits, including interpersonal reactivity index (IRI) and social value orientation (SVO). IRI^60^ was a measure of dispositional empathy, while SVO^61^ measured one’s prosocial and individualistic preferences about how to allocate resources (e.g., money) between the self and others. These two measures were selected to represent sociability capacity^62^. We hypothesized that the prosocial trait may modulate one’s self-serving motivation (e.g., caring more about others and less about self-interest) that drive dishonesty^7^. In line with our prediction, we found that the IRI scores were positively correlated with AUC (Pearson’s *r* = 0.49, *p* = 0.009, 95% CI = 0.14 to 0.73; Figure 3a, upper panel), indicating that it was more difficult for participants with a higher level of empathy to make a dishonest decision. Furthermore, the SVO score was negatively correlated with the total dishonesty rate (Pearson’s *r* = − 0.4, *p* = 0.037, 95% CI = -0.68 to -0.03; Figure 3b). Because we set higher rewards for the erroneous answers in most trials to induce dishonesty, and because dishonesty led to others’ loss of rewards, the more prosocial a participant was, the less likely they lied for a reward.

Next, we asked whether variation in personal traits affects the brain’s representation of dishonesty when considering reward and consistency. We conducted an IS-RSA to test which brain regions were responsible for representing the variation in personal traits. The IS-RSA procedure was similar to Figure 7a. The personal traits matrix was calculated based on IRI and SVO, and the brain matrix was based on the GLM beta map under two conditions in session 3. The correlation results indicated that participants with similar personal traits exhibit similar patterns in SMA, IPS, and SPG for the dishonesty condition in the last session (*FDR* correction, *p <* 0.05; Figure **??**a in *supplementary materials*). For the honesty condition in session 3, SMA, dACC, prACC, and PCC explained the inter-subject differences in personal traits (*FDR* correction, *p <* 0.05; Figure **??**b in *supplementary materials*). Together, the results showed that personal traits reflect variation not only in behavior but also in brain activity patterns.

### Choice entropy was linked to mouse tracking indices and related brain activities

We utilized entropy^63^ to quantify the overall randomness of participants’ choices. For every participants, we selected the menu with no timeout trial. For each item, four types of transition probability for responses (dishonesty/honesty) and reward difference (positive/negative) were conducted to calculated the choice entropy (for details, see *Methods*). While the difference of history responses reflected the trial-by-trial consistency, we averaged the choice entropy across items as a measure for the overall consistency. To examine the validity of this definition and the manipulation of our reward setting, we performed a linear mixed effect model, which turned out that reward difference entropy had a significant impact on choice entropy (*beta* = 0.18, *p* = 0.0002).

To investigate whether drift rates derived from trial-by-trial data had a relationship with the choice entropy, we found that choice entropy is positively correlated with the reward drift rate (Pearson’s *r* = 0.54, *p* = 0.0037, 95% CI = 0.20 to 0.76; Figure 9c). For mouse tracking indices, the choice entropy was positively correlated with the mouse tracking indices MAD and AD under honest responses in the last session (MAD: Pearson’s *r* = 0.42, *p* = 0.03, 95% CI = 0.044 to 0.69; AD: Pearson’s *r* = 0.47, *p* = 0.014, 95% CI = 0.11 0.72; Figure 9c and d). Besides, we found that the activities (dishonesty -honesty) of cognitive control and memory related regions in session 3 were also positively correlated with the choice entropy (PCC: Pearson’s *r* = 0.46, *p* = 0.016, 95% CI = 0.09 to 0.71; dACC: Pearson’s *r* = 0.59, *p* = 0.0012, 95% CI = 0.27 to 0.79; Hippocampus: Pearson’s *r* = 049, *p* = 0.0093, 95% CI = 0.14 to 0.73; Figure 9e).

## Discussion

Our study utilizes an experimental setting (Figure 2a) to test how people make decisions via the trade-off between reward and decision consistency. The variations in the trade-off of the two attributes were computationally characterized using a tDDM model that integrates previous history responses and reward magnitude. Further, we used IS-RSA to explore how variation in trade-offs (preference for reward and preference for consistency) mapped onto individual variations in brain activity and FCs over the brain regions of interest. Across individual and group analyses, the MT and fMRI results show that cognitive control and reward-related brain regions are involved in this reward and consistency trade-off and affect the hesitation when making decisions. Interestingly, the choice entropy is associated with indices of MT and the activity in the cognitive brain. Personal traits such as empathy also predict the outcome—people with higher levels of empathy showed less dishonesty. The findings show overall consistency and flexibility in moral decisions and the roles of reward and cognitive control brain in it.

### The roles of reward and consistency in the moral decisions

In the current binary decision task with an explicit presentation of the number of history responses and reward magnitude, we focused on the trade-off between maintaining consistency and seeking reward. The DDM results first suggest that both reward and consistency have significant impacts on moral decision making (Figure 4). Our finding that people weigh rewards more heavily in the dishonesty task resonates with studies that induce spontaneous immoral or dishonest behavior with monetary rewards as incentives^9,19^,^64,65^. This is reasonable because the desire for self-interest can result in lying, which may have negative effects on self-image. For example, the study shows that self-serving dishonesty gradually increases with repetition^19^. A recent fNIRS study shows how the reward system activates during the entire course of dishonest behavior and the way in which it affects dishonest decisions^66^. Our findings further suggest that such monetary reward-based manipulation is effective for motivated dishonesty^7,23^,^67^,^68^.

Although the reward plays a dominant role in moral decision-making, our results suggest that participants with a history of honest responses were less likely to lie, even when the reward for dishonest choices was higher (Figure 2). We set the dishonesty threshold as positive and the honesty threshold as negative. Therefore, these combined results indicated that the rewards drive participants to make dishonest decisions, whereas response history results in honest decisions. This finding reflects that consistency is an important factor to modulate the dishonesty rate. It is consistent with previous studies on self-consistency^13^, Decision history is an inescapable part of every similar or repeated decision, and heuristics offer individuals a general guide to follow^69,70^. This is important because it suggests that groups are more likely to make honest decisions when there is a conflict between honest response history and the reward offered. Furthermore, the tDDM shows that consistency drives honesty: although over 80% of trials were designed to induce dishonesty, a considerable number (about 50%) of participants only lied in approximately 50% of the trials (see Figure **??**). This study’s findings reveal an alternative explanation that despite a reward’s relatively stronger strength, the reward does not enter the decision process sufficiently early to influence choice. Combined with a previous study showing that children persevere more and make less strategic adjustments than adults^71^, the weight on choice history may be associated with the cognitive control system^72^. The previous decision reduces the effort in making the same decisions and leads to a gut decision. Such evidence also supports the view that consistently practicing honesty in some contexts enhances trust and honesty^72^.

### How are reward and consistency considerations associated with the cognitive control-related and reward-related brain regions

The cognitive control and reward brain systems are two interconnected neural networks that play crucial roles in moral decision-making^12,23^. They compete so that higher cognitive control may inhibit reward-seeking^73^. In this study, we tested two considerations regarding moral decisions, the amount of self-interest and self-consistency, under the framework of the interplay between reward and cognitive control brain. Given previous work indicating that the stability-flexibility balance in decisions is linked to cognitive control^74,75^, we provide evidence that the self-consistency process is associated with the cognitive control brain. Specifically, we provide evidence about how weights on reward and consistency are associated with cognitive control-and reward-related brain regions. First, we identified that the activity in the SMA is correlated with the weights of both consistency and reward, and SMA tracks reward and their interaction earlier than it tracks consistency. Consistent with earlier studies linking self-consistency and cognitive-control^76^–^81^, we observed that cognitive-control-related regions, such as vmPFC, dlPFC, prACC, and dACC, were directly correlated with the weights of consistency. Further, these brain regions mainly tracked consistency temporally (Figure 6). Regarding the reward-related brain region, the relationship between the weight of reward and the reward system was modulated by the mouse-tracking index AUC. The AUC reflects hesitation or the conflict present between two response options in the task^82^. Considering that self-interest can result in dishonesty, the AUC may reflect the degree of one’s hesitation to behave dishonestly, which is linked to the stronger or lower reward brain activity.

We found further evidence supporting our view of brain in reward and self-consistency trade-offs. In dishonesty responses, participants with similar evaluations of consistency and reward had similar activation patterns in the SMA, dlPFC, mPFC, ANG, IFG, and OTC. For the honesty condition, participants who similarly evaluated consistency and reward exhibited similar activation patterns in the pre-SMA, IFG, prACC, SMA, MTG, and STG (Figure 8). Combined with findings showing that the SMA and IFG are involved in proactive switching^83^,^84^, the findings suggest that the individual variations on the trade-off between consistency and reward indeed manifest in cognitive-control brain activity. Previous studies provide evidence on the role of the frontoparietal network (FPN)^85^–^90^ in switching behavior flexibly and adapting to changing environments. To examine the FC of reward and cognitive control-related brain with consideration of participants’ heterogeneity, dyadic regression was performed for the functional connectivity of prACC and ventral striatum with dlPFC, respectively. Studies show that the representation of rewards by individuals can predict dishonesty^22^,^91^. For example, the stronger neural activity or enhanced functional connectivity of the dorsolateral prefrontal cortex (dlPFC), ventromedial prefrontal cortex (vmPFC), anterior cingulate cortex (ACC), ventral tegmental area (VTA), and ventral striatum (VS) exhibited adjustments and described human sensitivity to rewards, and they can predict whether people engage in dishonest behavior. This study’s findings show that only consistency plays a significant role in predicting the connection between the dlPFC and prACC, whereas the reward plays a significant role in the connection between the dlPFC and VS. As such, we further validated our modeling results and uncovered the way in which consistency and reward interacted with the cognitive control and reward systems.

### Decision flexibility links to mouse trajectories, and the cognitive-control related brain regions

Notably, keeping self-consistency in decisions may require the interaction between cognitive control and the memory system. People use cognitive control to manage demands associated with memory^92^,^93^ of past response history and current self-interest temptation. Tracking past response patterns may help to weigh the contribution of consistency to decisions, and mechanisms of decision memory in moral decisions^94^. Here we propose another definition of consistency to quantify the overall randomness of choice order, which is the choice entropy^63^. The choice entropy was found to be correlated with the activity (dishonesty-honesty) of PCC, dACC and hipocampus in the last session, confirming the interaction between cognitive control related regions and memory system when keeping self-consistency. Also, it was correlated with mouse tracking indices MAD and AD, which measures the conflict between two choices. Further, to link the trial-by-trial consistency and the overall consistency, we found that the participant with higher reward drift rate had higher level of choice entropy, supporting the role of reward system in keeping self-consistency. Together, our results of choice entropy suggest the participation and interaction of cognitive control, memory and reward system in the pursuit of self-consistency.

Different from the above-described adaptions to dishonesty^19^, the current results show increased activation in the cognitive control-related brain regions throughout the sessions, especially in session 3. Interestingly, the significant increases in brain activation from the first to the last session mainly occurred in the honesty condition (Figure 5). One possible explanation is that most of the trials were designed to induce dishonesty, but the rate of lying might be lower during the first session. With the accumulation of history responses and the reward temptation, maintaining honesty in the later sessions might be more conflicting and require more control capacity. In this case, extra effort is required to switch responses and maintain honesty. The behavioral results are in line with the dishonesty adaption effect, which showed that reaction times decreased significantly along the sessions^19^. Additionally, the longer reaction times in providing dishonest responses echoed the Grace hypothesis that dishonesty requires extra cognitive resources^7^.

### how mouse tracking technique help reveal the decision dynamics

One novel aspect of this study is that we incorporated mouse tracking into the fMRI scanner, following previous studies^42^,^95^. Using these dynamic MT indices (e.g., MAD, AD, and AUC) reflecting conflicts and hesitation^7^,^96^, we obtained evidence on mental process associated with the trade-off between consistency and reward in three respects. First, on the behavioral level, we identified a significant bifurcation of mouse trajectories under the dishonesty and honesty conditions in the last session (Figure 3 right panel). Moreover, mouse measures, MAD, AD, and AUC presented a significant differentiation in the last session (Figure 9b in *Supplementary materials*). These results offered possible predictors for discriminating repeated dishonesty from honesty in future studies. Second, AUC modulated the relationship between the weight of reward and the reward brain (Figure 6c). Third, larger AUC in the last session is correlated with a higher IRI score (Figure 9a in *Supplementary materials*), indicating that participants with more empathy have more difficulty making a choice. These results align with those of previous studies suggesting social norms of personal values^7^,^67^. Mentalizing abilities^97^,^98^ were also related to dishonesty. Together, the MT results benefited this study by showing the differentiation and correlation of moral behavior and the brain in many aspects. Overall, the moral decision entails cognitive control and the reward brain system. Cognitive control regions enable individuals to consider potential consequences of self-image and control their actions, while the reward brain system can provide us with incentives to act in certain ways and guide our decision-making processes. However, the pursuit of rewards can interfere with moral norms, thereby highlighting the complex and dynamic nature of these decisions^94^. One potential limitation of the current study is the increased activation in motor cortex activity due to mouse tracking setup in the scanner, which may affect the RTs and DDM fitting even though our study focuses more on the social brain. Future studies may refine setups and provide new evidence on other decision-making tasks to fully understand the neural mechanisms underlying moral decisions and the theoretical frameworks of the reward-consistency trade-off.

Although the absence of self-consistency does not necessarily indicate moral hypocrisy, our results indicate it is still a factor for moral decision^8^. Self-consistency contributes to moral decisions with a link to cognitive control brain regions such as dlPFC, vmPFC, and prACC. The individual differences in moral trade-offs and fMRI signals further highlight the interaction of the cognitive control and reward brain and potentially can inspire future studies on morality training processes. Extending this view, a deeper understanding of decisions affected by external reward and internal consistency may help in the design of new types of AI ethic policy. Indeed, an early report has shown that chatGPT can provide inconsistent arguments on moral dilemmas, and human moral judgment is biased by ChatGPT^99^. Indeed, governments and scientists may need effective intervention in the ethic of AI and a better understanding of how external and internal factors influence human moral behaviors.

## Methods

### Participants

All experimental procedures were approved by the Institutional Review Board of the University of Macao (BSERE21-APP005-ICI). Regarding human subject recruitment, participant eligibility was first examined using an online pre-screening survey and determined by the experimenter using another detailed subject screening form. Exclusion criteria included claustrophobia, probability of pregnancy, history of heart disease and brain trauma, and the existence of any metal implant in body. To achieve greater than 80% power to detect a large effect of *d* = 0.80 at *α* = 0.05 in our analyses of paired t-tests (dishonesty and honesty condition), we calculated the minimal sample size, which was 14, using the package *pwr* in R^100^. We oversampled and recruited 37 participants via online advertisement from the University of Macao for the task. Two subjects were excluded due to missing behavioral data. Four subjects were excluded because they had a zero dishonesty rate. Four subjects were excluded to ensure that every participant lied in every session. Therefore, the sample size was 27 (13 females, mean age 20.89, *SD* = 2.75). The overall lying rate of the remaining subjects is presented in Figure S1 in the Supplementary Material. Participants were right-handed with normal or corrected-to-normal vision and had not participated in any similar studies. Each participant signed an informed consent form prior to the formal experiment, and the experimental protocol was approved by the local ethical review committee. At the end of the study, the participants were paid 130–150 MOP.

### Experimental Procedure

Stimuli were presented by Psychopy 2.3^101^ standalone. Participants viewed a 17-inch MR-ready LCD monitor (resolution: 2560 × 1600, refresh rate: 60 Hz), at 40 cm distance from the participants’ eyes at the end of the bore, via a helmet-mounted front-silvered mirror. A MRI-compatible optical mouse was used for participants to do the task in scanner. Meanwhile, during all functional sessions, mouse trajectory was recorded using PsychoPy 2.3^101^, sampled at 60 Hz.

The experiment was a self-paced task with nine runs, and each run consisted of the same set of 20 uncommon questions (details in *Supplementary materials*)) in a randomized order. As shown in Fig. 2a, the question was presented in the middle of the screen, and the start button was located at the bottom in the beginning of every trial. Participants were free to press the start button when they were prepared. Once they pressed the start button, two choices (one for the correct answer, the other for the incorrect one) were shown in the left and right corners of the screen. In addition to the correct and incorrect answers, other information for each choice was provided, including which was the correct answer (marked with a black circle), the monetary reward, and the value of how many times the option had been chosen for the same question item (marked as red triangles and would be shown in 2 − 9 runs). To induce participants to choose between reward and honesty, we set more money for the incorrect answer in over 60% of the trials. Participants had 4 s to answer by moving the computer mouse, and feedback was shown after their choice and lasted for 1 sec. If participants failed to provide valid responses within 4s, warnings were displayed, and that trial was excluded from further analyses. After participants finished one session, the cumulative monetary reward in the session was displayed on the screen, and they were able to take a rest until they were ready to continue. Crucially, once participants started the trial, the position of the mouse was automatically initiated at the middle bottom of the screen.

Before the formal task in the scanner, participants went through a training phase, which aimed to familiarize them with the experimental procedure. To avoid a situation in which participants only seek money and ignored the importance of being honest, they were told that this was a knowledge transmission task wherein the option they chose would be the preference for the latter participants to win the money. To ensure incentive compatibility, we set a greater reward for the error answer in 150 of 180 trials. Moreover, at the end of the experiment, the payments were calculated as the cumulative reward in the task. This procedure encouraged participants to treat each trial as if it may be one that would count for their monetary payment. After the formal task, participants completed the questionnaire, which consisted of the IRI^60^ and SVO^61^. The IRI scale is widely used for measuring individual differences in trait empathy and consists of four domains: (1) perspective taking (PT), (2) fantasy (FS), (3) empathic concern (EC), and (4) personal distress (PD). SVO is used to measure one’s individual preferences during social interactions^102^–^104^.

### Mouse Tracking Analysis

#### Mouse Trajectory Preprocessing

We extracted mouse trajectories from each session for each participant. Standard mouse-tracking preprocessing was conducted temporally and spatially^49^,^96^. In a typical binary choice design, trajectories end at either the left or the right response option. As the overall spatial direction is irrelevant for most analyses, all trajectories were remapped so that they would end on the same side. We used the R package ‘mousetrap’^105^ to map the trajectories to the left by default, suggesting that trajectories that end on the right-hand side are flipped from right to left. We rescaled all mouse trajectories into a standard coordinate space (top left: [-1, 1]; top right: [1,1]) so that the cursor always started at [0,0]^96^,^106^. Temporally, time normalization was applied to the trajectories such that the duration of each trial was divided into 101 identical time bins using linear interpolation to obtain the average of their length across multiple trials^35^,106–^110^.

#### Mouse Trajectory Measurements

Several different measures for the curvature of mouse trajectories have been proposed in the literature^49^,^111^. One frequently used measure is the MAD. The MAD represents the maximum perpendicular deviation of the actual trajectory from the idealized trajectory, which is the straight line connecting the trajectories’ start and end points. The MAD and many additional trial-level measures can be calculated using the mtmeasures function. These measures are then aggregated per participant for each level of the within-participants factor. Another frequently used measure is AUC. AUC represents the geometric area from the reference trajectory to the actual trajectory. The higher the AUC is, the bigger the attraction to the other response option. AUC is a good indicator of the overall attraction^49^. The third index we calculated was the AD, which is sensitive to the temporal dynamics of the movement^112^.

#### Mouse Trajectory Comparison

We averaged all mouse trajectories into two conditions (dishonest and honest) of each sliding window after preprocessing them. To test the significant temporal difference between the two conditions statistically, we calculated the positions of each time step in every trial for the two conditions and compared them through a paired t-test (fdr corrected)^105^,^113^. We used the results to infer in which time step the mouse trajectory began to bifurcate in each sliding window on an individual level^96^.

### tDDM modeling

All statistical analyses for the behavioral data were carried out in R^114^.

#### The tDDM

We utilized a multi-attribute tDDM), in which reward and consistency could affect the evidence accumulation process with different onset times and weights. Specifically, the evidence (*E* updated in 4-ms steps per convention, that is, *dt* = 4*ms*. At each step, a weighted amount of relative (error choice minus correct choice, that is, dishonest choice minus honest choice) reward (*MD*) and former choices (*CD*) was added to the evidence (as is shown in equation. 1). When the evidence reached the threshold (set as 2) for the error answer (dishonest; equals 2.5) or the correct answer (honest; equals -2.5), a decision was made for that answer.

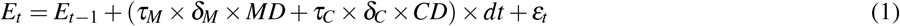

where

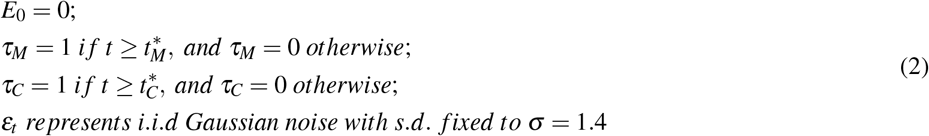

In this model, there are four free parameters to estimate separately for each participant. The initial value of the evidence (*E*_0_) was set as 0; *δ*_*M*_ and *δ*_*C*_ represented the drift rates as well as the weights for reward and consistency. 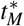 and 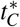 represented the onset times that reward and consistency came into the evidence accumulation process. Once the absolute value of evidence (*E*_*t*_) evolved to the threshold, a decision was made: if *E*_*t*_ reached the positive boundary, a dishonest choice was made; if *E*_*t*_ reached the negative boundary, a honest choice was made. The final reaction time was computed as *t* × *dt*. Moreover, this model assumed that the non-decision time proposed in standard DDMs was included in the time window before 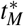 and 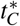, which accounted for the stage required for any initial perceptual or subsequent motor processes that surround the period of active evidence accumulation and comparison.

#### Alternative DDMs

We also fitted five alternative DDM formulations to our data. mDDM(multi-attribute DDM) assumed that two attributes (reward and consistency) started influencing the evidence accumulation process at the same onset time but with different drift rates (where 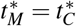, *τ*_*M*_ = *τ*_*C*_). latDDM (latency DDM) assumed that onset times but not drift rates varied by attribute (where *δ*_*M*_ = *δ*_*C*_). sDDM (standard DDM or simple DDM) presumed reward and consistency shared the same drift rate and onset time (where 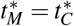, *τ*_*M*_ = *τ*_*C*_, *δ*_*M*_ = *δ*_*C*_) (See Equations in *Supplementary materials*).

#### Model fitting

The best values of the four parameters were estimated on the subject level (reward and consistency drift rates and onset times) for tDDM. We collapsed choices into a binary dishonesty/honesty choice, and reaction times of honesty trials were set as negative^115^. We used the differential evolution algorithm^116^ with 150 iterations for each participant. For each iteration, 3000 reaction times and decisions were simulated for each unique combination of reward and former choice values. According to the distribution of the simulated data, we computed the likelihood of our experimental data and maximized it in the subsequent iteration. By examining the value of the four parameters for every generation, we found that the best-fitting parameters were determined before 150 iterations. We fit the alternative DDMs to the empirical data using the same procedure.

#### Parameter recovery

The best-fitting parameters were used to perform parameter recovery tests to validate our model. We generated simulated choices and reaction times according to the best-fitting parameters of tDDM. The simulated choice sets were based on the consistency and reward differences participants faced during each decision trial. Thus, the simulated choice sets matched the empirical data in terms of trial numbers and attribute difference distributions. Fitting these simulated choices allowed us to quantify the ability of the best-fitting model to recover known parameter values as well as the ability to distinguish between these models.

### fMRI acquisition

All fMRI data were acquired using a 3.0 T Siemens MAGNETOM Prisma MRI scanner with a 64-channel head coil in the Center for Cognitive and Brain Sciences, University of Macau. Neuroimaging data acquisition included the collection of both anatomical and functional data. High-resolution T1-weighted images were acquired for each participant (3D MPRAGE sequence; voxel size = 1 mm isotropic; FOV = 256 mm; 176 slices, slice thickness: 1.0 mm; TR = 2300 ms, TE = 2.26 ms, TI = 900 ms, flip angle = 8°). Whole-brain functional imaging was acquired by a T2*-weighted gradient echo, echo-planar pulse sequence in descending interleaved order with a 2.0mm slice gap, voxel size of 2.0 mm, slice thickness of 2.0 mm, TE of 30 ms, TR of 1000 ms and flip angle of 90°.

### fMRI data preprocessing

The fMRI data preprocessing was based on Nipype^117^, and was implemented in fMRIPrep v20.2.1^118^ with the default pipeline.

#### Anatomical data preprocessing

A total of 1 T1-weighted (T1w) images were found within the input BIDS dataset. The T1-weighted (T1w) image was corrected for intensity non-uniformity (INU) with N4BiasFieldCorrection^119^, distributed with ANTs 2.3.3 [120, RRID:SCR 004757], and used as T1w-reference throughout the workflow. The T1w-reference was then skull-stripped with a *Nipype* implementation of the antsBrainExtraction.sh workflow (from ANTs), using OASIS30ANTs as target template. Brain tissue segmentation of cerebrospinal fluid (CSF), white-matter (WM) and gray-matter (GM) was performed on the brain-extracted T1w using fast(FSL 5.0.9, RRID:SCR 002823)^121^. Volume-based spatial normalization to one standard space (MNI152NLin2009cAsym) was performed through nonlinear registration with antsRegistration (ANTs 2.3.3), using brain-extracted versions of both T1w reference and the T1w template. The following template was selected for spatial normalization: *ICBM152 Nonlinear Asymmetrical template version 2009c*(RRID:SCR 008796; TemplateFlow ID: MNI152NLin2009cAsym)^122^.

#### Functional data preprocessing

For each of the 3 BOLD sessions found per subject (across all tasks and sessions), the following preprocessing was performed. First, a reference volume and its skull-stripped version were generated using a custom methodology of *fMRIPrep*. The BOLD reference was then co-registered to the T1w reference using flirt(FSL 5.0.9)^123^ with the boundary-based registration^124^ cost-function. Co-registration was configured with nine degrees of freedom to account for distortions remaining in the BOLD reference. Head-motion parameters with respect to the BOLD reference (transformation matrices, and six corresponding rotation and translation parameters) are estimated before any spatiotemporal filtering using mcflirt (FSL5.0.9)^125^. BOLD sessions were slice-time corrected using 3dTshift from AFNI 20160207 (RRID:SCR 005927)^126^. The BOLD time-series were resampled onto their original, native space by applying the transforms to correct for head-motion. These resampled BOLD time-series will be referred to as *preprocessed BOLD in original space*, or just *preprocessed BOLD*. The BOLD time-series were resampled into standard space, generating a preprocessed BOLD session in *MNI152NLin2009cAsym space*. First, a reference volume and its skull-stripped version were generated using a custom methodology of *fMRIPrep*. Several confounding time-series were calculated based on the preprocessed BOLD: framewise displacement (FD), DVARS and three region-wise global signals. FD and DVARS are calculated for each functional session, both using their implementations in *Nipype* (following the definitions by^127^). The three global signals are extracted within the CSF, the WM, and the whole-brain masks. Additionally, a set of physiological regressors were extracted to allow for component-based noise correction (*CompCor*)^128^. Principal components are estimated after high-pass filtering the *preprocessed BOLD* time-series (using a discrete cosine filter with 128s cut-off) for the two *CompCor* variants: temporal (tCompCor) and anatomical (aCompCor). tCompCor components are then calculated from the top 2% variable voxels within the brain mask. For aCompCor, three probabilistic masks (CSF, WM and combined CSF+WM) are generated in anatomical space. The implementation differs from that of Behzadi et al. in that instead of eroding the masks by 2 pixels on BOLD space, the aCompCor masks are subtracted a mask of pixels that likely contain a volume fraction of GM. This mask is obtained by thresholding the corresponding partial volume map at 0.05, and it ensures components are not extracted from voxels containing a minimal fraction of GM. Finally, these masks are resampled into BOLD space and binarized by thresholding at 0.99 (as in the original implementation). Components are also calculated separately within the WM and CSF masks. For each CompCor decomposition, the *k* components with the largest singular values are retained, such that the retained components’ time series are sufficient to explain 50 percent of variance across the nuisance mask (CSF, WM, combined, or temporal). The remaining components are dropped from consideration. The head-motion estimates calculated in the correction step were also placed within the corresponding confounds file. The confound time series derived from head motion estimates and global signals were expanded with the inclusion of temporal derivatives and quadratic terms for each^129^. Frames that exceeded a threshold of 0.5 mm FD or 1.5 standardised DVARS were annotated as motion outliers. All resamplings can be performed with *a single interpolation step* by composing all the pertinent transformations (i.e. head-motion transform matrices, susceptibility distortion correction when available, and co-registrations to anatomical and output spaces). Gridded (volumetric) resamplings were performed using antsApplyTransforms (ANTs), configured with Lanczos interpolation to minimize the smoothing effects of other kernels^130^. Non-gridded (surface) resamplings were performed using mri vol2surf (FreeSurfer).

### fMRI GLM analysis

We performed a standard first-level GLM approach with three scan sessions (nine runs). A GLM was constructed for each participant using decisions as regressors. The time at which participants clicked on the start button was treated as the onset time. The reaction time of each trial was added as both parametric and temporal modulators to account for the height and duration of the BOLD signal, respectively^47^.

The resulting GLM was convolved with SPM’s canonical hemodynamic response function. The model was corrected for temporal autocorrelations using a first-order autoregressive model, and a standard high-pass filter (cutoff at 128 s) was used to exclude low-frequency drifts.

We obtained the beta maps of two decision types (dishonesty or honesty) and their contrast in each session. The beta maps estimated from the first-level GLM were used in all subsequent analyses.

### ROI selection and beta extraction

According to the previous studies, keeping a self-image involves cognitive-control related brain regions like dlPFC and ACC. Reward involves ventral striatum and OFC. we selected 5 ROIs for consistency including SMA, prACC, dACC, dlPFC and vmPFC, and 2 ROIs for reward including ventral striatum and OFC (see Fig 6b. Specifically, masks of SMA, dlPFC, vmPFC and ventral striatum(VS) were obtained from meta-analysis from Neurosynth website (https://neurosynth.org/). Others were created via Brodmann atlas by using WFU PickAtlas toolbox^131^.

Beta maps within the ROIs from the first level GLM were extracted using nltools^132^.

### Single-trial beta time courses

We used each ROI mask to extracted preprocessed BOLD time courses as the average of voxels within the mask. For each scan session, we regressed out variation due to head motion acquired from fMRIPrep, applied a high-pass filter (128 s cut-off) to remove low-frequency drifts, and oversampled the BOLD time course by a factor of 23 (time resolution of 0.144 s; spline interpolation). For each trial, we extracted activity estimates in a 8 s window after the trial onset^133^.

Then to acquire the beta time series for two attributes (reward and consistency) respectively, we performed two GLMs on subject level. we regressed the time courses over several regressors. For ROIs that had correlation with the tDDM parameter under both two conditions like SMA and vmPFC (see results), we added trial-wise consistency differences, reward difference and the interaction of these two as regressors. For reward-related ROIs, we perform the same procedure. For other ROIs, the interaction of response (dishonesty = 1, honesty = 0) and consistency difference, the interaction of response and reward difference and the interaction of reward and consistency difference were treated as regressors. We regressed the preprocessed time courses on these regressors for each time points, and plotted the coefficient time courses.

### Inter-subject representational similarity analysis (IS-RSA)

We used intersubject representational similarity analysis (IS-RSA)^134^,135 in consideration of tDDM parameters to identify regions of the brain that were responsible for reward and consistency. We mainly focused on session 3 because in the last session participants have fully taken history choices into account than the first two sessions. The IS-RSA analysis was performed in Python 3.9 using the NLTools package version 0.4.7 (http://github.com/ljchang/nltools).

We first obtained each participant’s activity map in the univariate analysis by extracting the GLM beta maps for the certain condition of session 3. We then divided these subject-level beta maps into 200 parcels using a whole-brain parcellation based on meta-analytic functional coactivation of the Neurosynth database (parcellation available at http://neurovault.org/images/39711/ and displayed in Supplementary Figure **??**). This parcellation scheme was chosen because of its computationally economization and non-overlapping parcellation over the whole cortex^135^.

Next, inter subject pairwise correlation for each response type (dishonest or honest) was calculated by *Scipy*’s Spatial module to carry out a dissimilarity matrix for each parcel (the “parcel dissimilarity matrices”). For tDDM’s parameter space, we also created an inter subject dissimilarity matrix using the Euclidean distance by using the drift rates of reward and consistency.

Last, we computed the Spearman’s rank-order correlations between each condensed parcel dissimilarity matrix and the condensed model dissimilarity matrix. This explores brain regions encoding the reward and consistency evaluation. To obtain significance levels of the resulting Spearman’s *rhos*, we performed the permutation tests by shuffling the order of the observations in model dissimilarity matrix 10,000 times, and calculated the proportion of the permuted *rhos* that exceeded the true *rho*. Lastly, these *p*-values were *FDR*-corrected and the threshold was set at 0.05.

### Functional connectivity analysis

To explore how the value signal in the dlPFC interacts with consistency-related (dACC) and reward-related (ventral striatum, VS) brain during the trade-off process, we implemented a functional connectivity analysis using the CONN toolbox^59^ (version 21.a) (https://www.nitrc.org/projects/conn) for session 3.

We employed a standard pipeline by using the pre-processed fMRI data. In the denoising step we used linear regression to remove the influence of the following confounding effects on the fMRI time course: (1) BOLD signal from the white matter and CSF voxels (five components each, derived using the anatomical component-based correction (aCompCor) implemented using the ART toolbox), (2) eighteen confounds by fMRIPrep: six residual head motion parameters, global signal (the average signal within the brain mask), framewise displacement (the quantification of the estimated bulk-head motion calculated using formula proposed by^136^), six additional noise components calculated using anatomical CompCor and four DCT-basis regressors, (3) effect of task-condition using separate run regressors (for dishonesty and honesty conditions) convolved with the haemodynamic response function. Thus, we performed the connectivity analysis on the residuals of the BOLD time series after removing condition-related activation/deactivation effects^59^,^137^,^138^. Finally, the denoising step included temporal bandpass filtering (0.008–0.09 Hz), and linear detrending of the functional time course. Following pre-processing, we performed condition-dependent functional connectivity analysis on the mean BOLD time course^59^ extracted from selected ROIs.

### Representational dissimilarity matrix GLM

To investigate how functional connectivity calculated above was associated with DDM parameters and personal traits, we constructed inter-subject representational dissimilarity matrix (RDM) of functional connectivity (dlPFC-dACC and dlPFC-ventral striatum), drift rate of reward, drift rate of consistency, IRI score and SVO score. Because these matrices had different scales, we calculated the standardized euclidean distance between subjects by *scipy*’s Spatial module. Respectively, we regressed two functional connectivity over the rest four RDMs (drift rate of consistency, drift rate of reward, IRI score and SVO score) by using *lm* function in R.

### Entropy of responses

Entropy^63^ is one way to quantify the randomness of a system. After excluding the question with timeout trials, we selected responses of 9 runs for the last questions. Transitions of responses for a certain question were categorized into 4 types: dishonesty -honesty, dishonesty -dishonesty, honesty -dishonesty, and honesty -honesty. Thus we formalized a transition 2-by-2 matrix N where the rows represent the current response and the columns represent the next response, with the probability of transition in each cell:

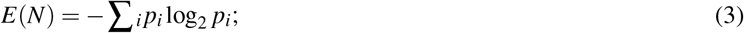

where i represented the type of transitions. We calculated the entropy for each question and averaged on subject level.

## Data and code availability

All data reported in this paper will be shared by the lead contact upon request. All original code has been deposited on GitHub: https://github.com/andlab-um/RDdishonesty and is publicly available as of the date of publication. Any additional information required to reanalyze the data reported in this paper is available from the lead contact upon request.

## Acknowledgements

This work was mainly supported by the Natural Science Foundation of China (U1736125), Science and Technology Development Fund (FDCT) of Macau [0127/2020/A3, 0041/2022/A], the Natural Science Foundation of Guangdong Province (2021A1515012509), MYRG of University of Macau (MYRG2022-00188-ICI), Shenzhen-Hong Kong-Macao Science and Technology Innovation Project (Category C) (SGDX2020110309280100), and the SRG of University of Macau (SRG2020-00027-ICI). We would like to thank Prof. Daeyeol Lee for helpful comments on the early version of the manuscript. We would like to thank all participants that took part in the study and enabled this research to be possible.

## CRediT author statement

Xinyi Julia Xu: Conceptualization, Methodology, Software, Data curation, Formal analysis, Visualization, Writing – review and editing. Guochun Yang: Methodology, Writing – review and editing. Jiamin Huang: Data curation. Ruien Wang: Data curation. Haiyan Wu: Conceptualization, Methodology, Software, Data curation, Formal analysis, Visualization, Writing – review, editing, Supervision, and Funding acquisition.

## Footnotes

1 Our sample initially constituted 37 participants, but we excluded 10 participants to ensure that every participant lied in every session (see detailed exclusion criteria in the Methods section). Data obtained from 27 participants were reported in the results.

